# SenoQuant: One-stop AI software for senescence marker analysis and prediction

**DOI:** 10.64898/2026.06.18.733222

**Authors:** Anthony B. Lagnado, Yukang Li, Chibuzo C. Nwakama, Ana Catarina Franco, Yeaeun Han, Diana Jurk, Helene Martini, Stella Victorelli, Gung Lee, Dominik Saul, Adam J. Hruby, Lilian Salez Gomez, Seunghwa Woo, Joshua Farr, Saranya Wyles, Latifa Khalfaoui, Daniela G. Costa, Miiko Sokka, Sundeep Khosla, Nicola Neretti, YS Prakash, Jon Camp, David Holmes, João F. Passos

## Abstract

Senescent cells accumulate with age and contribute to tissue dysfunction, yet their identification in tissues is challenging due to low abundance, heterogeneous phenotypes, and the lack of specific markers. Senescence-associated features span multiple subcellular compartments, including nuclear DNA damage foci, cytosolic protein changes, and perinuclear alterations, each requiring tailored detection strategies. To overcome these challenges, we developed SenoQuant (https://github.com/HaamsRee/senoquant), a versatile software designed for comprehensive, accurate, and unbiased spatial quantification and prediction of senescence markers across diverse tissue contexts. Utilizing AI models, SenoQuant enables precise nuclear and cytoplasmic segmentation and detection of senescence markers across low- and high-plex imaging modalities, applicable to cultured cells and tissue sections from mice and humans. The platform also supports custom AI models; for example, we built SenCeption, a proof-of-concept predictor of single-cell p21 status from DAPI-stained nuclei in human skin. Available as a free napari plugin, SenoQuant is widely accessible to researchers. By providing a unified approach to senescence analysis and prediction, SenoQuant opens new opportunities for exploring the complex biology of senescence and its impacts on aging and disease.

## Introduction

Cellular senescence is an irreversible growth arrest state induced by various stressors, characterized by the secretion of pro-inflammatory cytokines, chemokines, and matrix-degrading proteins, collectively known as the senescence-associated secretory phenotype (SASP)^1^. These cells accumulate with age in different tissues and contribute to the pathogenesis of several chronic diseases^2^. Senescent cells influence multiple aspects of cell biology, including chromatin remodeling, organelle function, and SASP secretion^3,4^. However, identifying senescent cells remains challenging due to the lack of a definitive marker. Commonly used indicators like SA-β-Gal activity or elevated p16 and p21 levels often lack specificity, as these features can be found in non-senescent cells under certain physiological conditions^3,4^.

Recent advances in single-cell technologies have enabled better mapping of senescent cells, yet these methods frequently overlook spatial information. Spatial context is crucial, as senescent cells interact with surrounding cells, affecting their function and recruiting immune cells through SASP secretion^5-8^. This highlights the urgent need for spatially resolved methods that can quantitatively map senescent cells and their interactions with the immune system^9^.

Current spatially-resolved imaging techniques face significant challenges in extracting reliable information about senescent cells^9^. The complexity of senescence-associated markers demands distinct analytical techniques. For instance, markers such as DNA damage foci^10^, telomere-associated foci^11,12^, or centromere decondensation^13^ require precise detection of nuclear foci and/or colocalization in three-dimensional images. Similarly, methods like RNA *in situ* hybridization which are commonly used to measure senescence-associated markers, must accurately quantify transcripts within the nucleus or cytoplasm^14,15^. The analysis of lamin B1, which decreases in senescent cells, involves precise fluorescence intensity measurements around the perinuclear region^16^. Markers such as SASP expression or β-Gal activity necessitate quantification of cytoplasmic fluorescence intensity^17^. Moreover, given that senescence-associated markers may be present at very low frequencies^18^, accurate cell segmentation adaptable to varying tissue densities is crucial. Finally, recognizing that no single marker can reliably identify senescent cells, it has become essential to simultaneously analyze multiple markers within the same tissue section using spatial omics techniques^9^. However, manual quantification of spatial omics datasets is labor-intensive, time-consuming, and prone to biases. As part of the NIH-Common Fund’s SenNet initiative^19^, which aims to measure senescence across multiple tissues during aging using a range of methodologies, we identified the absence of a comprehensive tool to unify and integrate these diverse analytical approaches^9^.

To facilitate the analysis of the complex image modalities used to study cellular senescence, we developed SenoQuant (https://github.com/HaamsRee/senoquant), a versatile software designed to facilitate the measurement and integration of multiple imaging datasets. SenoQuant integrates the napari (https://github.com/napari/napari) image viewer and user interface to provide a no-code and user-friendly analysis environment. It features 2D and 3D deep learning models^20-22^ and morphological operations fine-tuned for accurate nuclear and cytoplasmic segmentation in a wide range of murine and human tissues. Additionally, to encourage the development and adoption of more capable Artificial Intelligence (AI) models to predict senescence *in vivo*, SenoQuant integrates a standard interface that allows custom models to make predictions on senescence-associated biomarkers from imaging-based features.

Taking advantage of SenoQuant’s model interface, we developed SenCeption, an AI classifier that accurately identifies individual p21-positive cells in human skin from DAPI alone. SenCeption is a convolutional neural network (CNN) that uses features available through DAPI staining as inputs and predicts whether the cell at its input is p21-positive or p21-negative. It achieves 80% accuracy in an independent testing dataset consisting of PhenoCycler-imaged skin sections. To our knowledge, this is the first classifier of its kind that achieves single-cell accuracy *in vivo*. Our results demonstrate the feasibility of adapting AI models designed for computer vision in predicting senescence-associated features *in vivo* at single-cell resolution with minimal input information.

SenoQuant enables comprehensive analysis and prediction of senescence-associated markers across diverse tissue contexts, providing a unified tool for generating accurate, unbiased, and spatially resolved insights into multiple features of senescence. Additionally, SenoQuant is available as a downloadable plugin within the free software napari, making it broadly accessible to researchers. Here, we demonstrate how SenoQuant measures multiple senescence-associated markers using both low- and high-plex methods, with applications ranging from cell cultures to tissue sections in mice and humans. We also show how DAPI staining and nuclear segmentation masks, both available at SenoQuant’s AI prediction interface, can be used to develop SenCeption. By integrating advanced image analysis capabilities with user-friendly workflows, SenoQuant significantly reduces the time required for quantification while also minimizing potential biases associated with manual measurements. Its ability to detect and quantify multiple senescence-associated markers across diverse tissue types and imaging formats offers a comprehensive approach to studying senescence. Meanwhile, SenCeption represents the next step in senescence detection *in vivo* and demonstrates the feasibility of using detailed tissue-derived data to develop accurate single-cell predictors.

## Results

### Overview of SenoQuant’s core features

To use SenoQuant, users first install the software using the provided installer (available for most operating systems) and launch it from the desktop icon. SenoQuant supports a wide range of microscopy file formats (e.g., lif, tiff, nd2, czi, ome-tiff, ome-zarr, etc.). Images can be opened either by drag-and-drop or via File > Open File(s). Additionally, the user can search for and download available SenNet spatial proteomics datasets through the integrated SenNet portal, enabling further analysis. Once an image is opened, channels are automatically separated and named.

The GUI (**Extended Data Figure 1**) follows a step-by-step workflow, allowing parameter setting and result validation at each stage. The main workflow includes (**Figure 1a**):

**1) SenNet portal**: SenNet publishes a broad range of multi-omics datasets, including multiplexed spatial proteomics (https://data.sennetconsortium.org/). Search for and download available SenNet spatial proteomics datasets in a convenient interface.
**2) Segmentation**: Use deep learning models and morphological operations to accurately segment nuclei and cells.
**3) Spot detection**: Detect and segment spot-like signals in 2D and 3D.
**4) Prediction**: Integrate and run custom AI predictors to obtain senescence-associated feature maps.
**5) Quantification**: Quantify and export per-cell/nucleus/spot fluorescence marker intensity, morphology, spot count, colocalization, etc. to CSV or XLSX tables.
**6) Visualization**: Visualize exported data tables using multiple plot types, including spatial plots, UMAP projections, double-expression plots, and neighborhood enrichment analyses (**Extended Data Figure 2**).
**7) Batch processing**: Run the same settings across multiple images and obtain quantification results.
**8) Settings saving and loading**: Save and load batch settings and user interface states for reproducible analyses.

**Figure 1.**
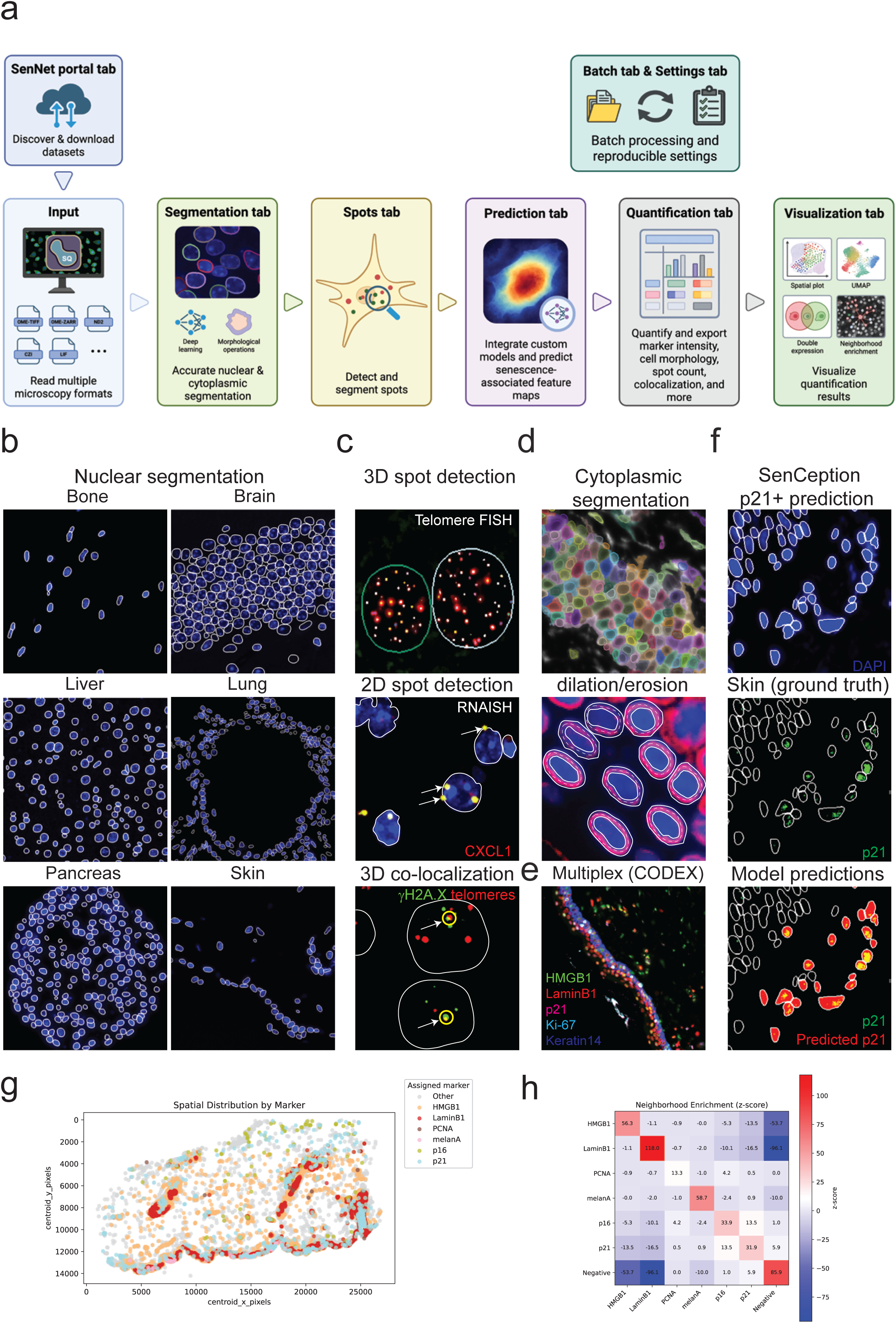
Summary of SenoQuant’s features and their applications in senescence research. **(a)** Schematic representation of the SenoQuant workflow. Following image acquisition, raw images are imported into SenoQuant. After reading the images and separating channels, each channel can be assigned to one of the following tabs: segmentation, spots, and prediction. Following segmentation (nuclear, cytoplasmic, or spots), the Quantification tab can be used to configure features and export results with any number and combination of channels and segmentations. The Visualization tab allows the user to generate various plots with the exported results. Settings can be saved and loaded in the Settings tab and also used in the Batch tab to process multiple images sequentially without user intervention; **(b)** Representative images of nuclear segmentation using SenoQuant’s default_3d model in bone, brain, liver, lung, pancreas, and skin (nuclear masks are overlaid in white on DAPI-stained nuclei in blue). **(c)** Representative images demonstrating SenoQuant’s spot detection capabilities: **(upper panel)** 3D spot detection applied to telomere detection using FISH (spot labels overlaid in multiple colors on the red telomere signal); **(middle panel)** 2D spot detection applied to CXCL1 RNA-ISH (yellow spot labels overlaid on the red RNA signal); **(lower panel)** 3D colocalization analysis applied to a telomere-associated foci (TAF) assay, showing spatial colocalization between telomeres (red) and γH2A.X (green) with yellow ring overlays; **(d)** Representative images of cell membrane segmentation (membranes overlaid in various colors) and nucleus dilation/erosion applied to lamin B1 detection (white double-membrane overlay on red lamin B1 signal); **(e)** Multiplex imaging application using PhenoCycler (formerly CODEX) in human skin. Nuclear segmentation (yellow) is overlaid on DAPI (blue), with additional markers including HMGB1 (green), lamin B1 (red), p21 (pink), Ki67 (light blue), and keratin 14 (dark blue); **(f)** Representative images of the AI prediction interface’s output when using the SenCeption model to predict p21; **(g), (h)** Representative plots generated by the Visualization tab. Spatial distribution by marker and neighboring enrichment analysis (z-score).

Users can easily navigate through the tab-based interface and perform each step independently.

SenoQuant’s development included building, fine-tuning, and integrating AI models for nuclear segmentation in 2D and 3D DAPI-stained tissues. These models were trained on publicly available datasets as well as manually annotated images generated in our laboratory. Validation showed that the models accurately identified nuclei in 2D and 3D images from various tissues, including bone, brain, liver, skin, pancreas, amongst others (**Figure 1b, Extended Data Figure 3**).

In addition, SenoQuant integrates deep learning and morphological spot detectors for segmentation of spot-like signals in 2D and 3D images. The currently implemented algorithms include rotational morphological processing (RMP)^23^ and U-FISH (https://github.com/UFISH-Team/U-FISH)^24^. U-FISH was modified to output spot segmentations rather than only spot coordinates, which is necessary for colocalization analysis. Both the RMP and U-FISH implementations include custom parameters and pre- and post-processing steps to improve performance. These algorithms are useful for detecting foci, such as nuclear telomere FISH, γH2A.X and RNA-*in situ* hybridization signals in both 2D and 3D images, and colocalization between spots, like telomeres and γH2A.X, which are used as markers of senescence (**Figure 1c**).

SenoQuant also enables cell membrane detection using two types of methods: (1) Deep learning (Cellpose-SAM)^22^ and (2) morphological operations (nuclear dilation and perinuclear rings)^25^. These methods allow quantification of cytosolic senescence markers. For example, the perinuclear rings algorithm is particularly effective for detecting lamin B1 in the perinuclear region (**Figure 1d**).

In the “Quantification” tab, nuclear, cytoplasmic, and spot-like features can be synthesized to output structured tables. SenoQuant supports exporting an unlimited number of segmentations and imaging channels for spatial omics applications (**Figure 1e**).

Moreover, SenoQuant’s AI prediction interface allows the user to integrate any custom models to generate senescence-associated feature maps. The outputs, if in image format, can then be saved and exported in Quantification (**Figure 1f**).

Finally, SenoQuant provides support to create various graphs using the exported data tables, such as spatial distribution by marker (using user-defined thresholds) and spatial neighborhood enrichment (z-score) (**Figure 1g,h, Extended Data Figure 2**).

### Detection of senescence-associated markers in cultured cells

To evaluate the capacity of SenoQuant to detect senescence-associated markers *in vitro*, we induced senescence in human MRC-5 fibroblasts using X-ray irradiation using established protocols^26^. We then assessed the expression of commonly used senescence-associated markers p21 and p16, as well as the absence of Ki67 and the loss of nuclear HMGB1, through immunofluorescence (**Figure 2a-e**). We proceeded to quantify the mean intensities of markers. We also utilized SenoQuant’s spot detector to quantify γH2A.X in senescent cells **(Figure 2f)**. The results demonstrated that SenoQuant is capable of accurate nuclear segmentation of proliferating and senescent fibroblasts grown *in vitro* and can be used to quantify the nuclear intensity of commonly used senescence-associated markers, showing significant differences between proliferating and senescent cells with similar values as reported previously by our laboratory and others^11,16,26,27^. Furthermore, it can be used to quantify nuclear diameter, which has been shown to increase in senescent cells^28^ (**Figure 2g**).

**Figure 2.**
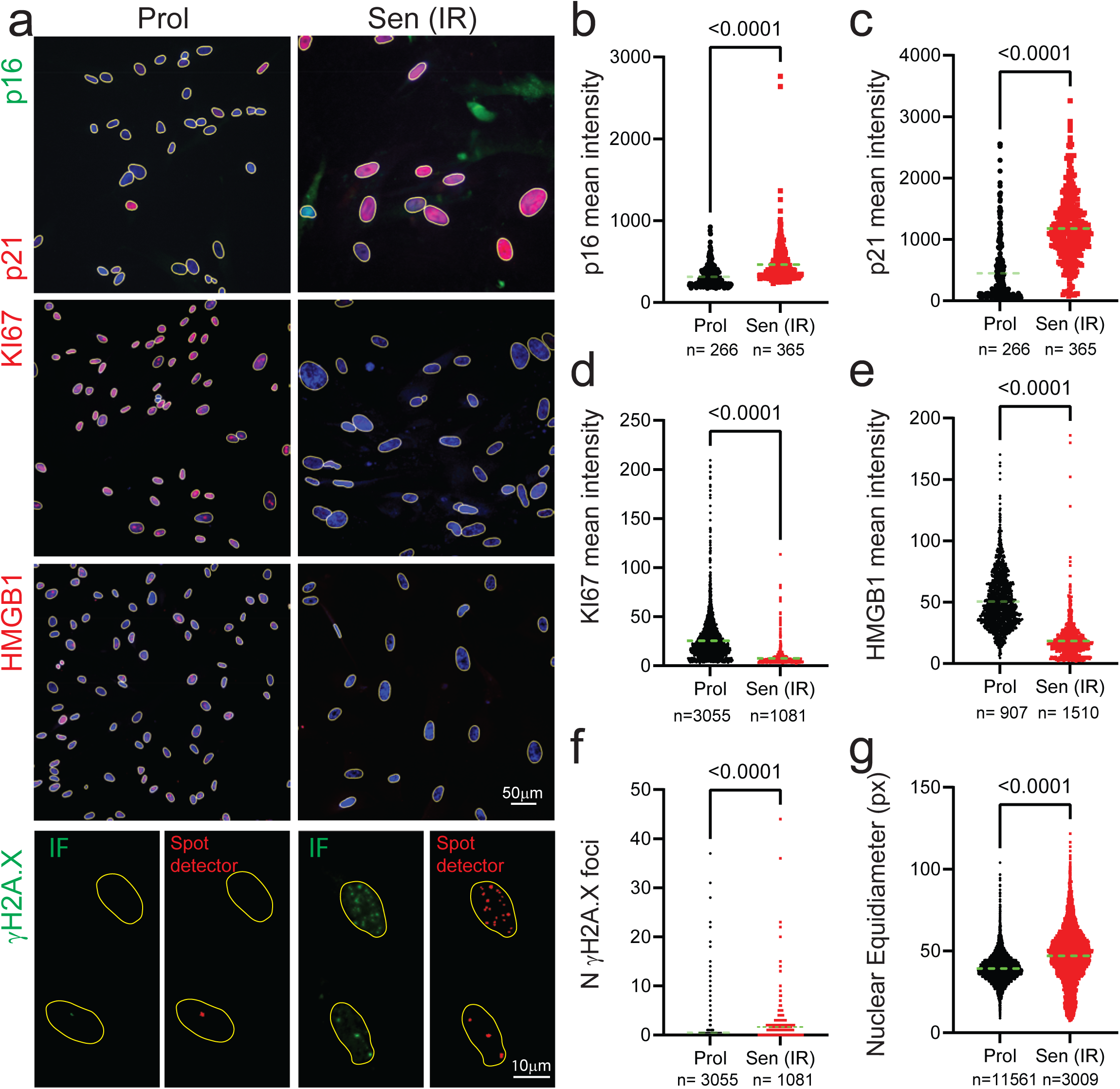
Using SenoQuant to quantify senescence-associated markers *in vitro*. **(a)** Representative images of proliferative (Prol) and senescent irradiated (Sen IR) MRC5 fibroblasts (10 days post-irradiation), showing nuclear segmentation overlays (yellow) alongside immunostaining for senescence-associated markers: p21 (red), p16 (green), Ki67 (red), HMGB1 (red), γH2A.X (green), and spot detector results (red); **(b-e)** Quantification of mean fluorescence intensity for nuclear markers using SenoQuant nuclear segmentation and marker channel assignment (process tab): **(b)** p16, **(c)** p21, **(d)** Ki67, and **(e)** HMGB1; **(f)** Quantification of mean γH2A.X foci per cell using SenoQuant’s spot detector and nuclear segmentation; **(g)** Measurement of mean nuclear equidiameter (in pixels) per cell using SenoQuant’s nuclear segmentation parameters. Scale bars: 50 µm (20× objective) and 10 µm (63× objective). Data are shown as dot plots with the mean indicated in green. The number of cells analyzed is shown below each graph. Results are from 3-4 independent experiments. Numerical p-values are indicated in each graph (Mann-Whitney).

### Quantifying perinuclear lamin B1 intensity using SenoQuant’s nuclear dilation feature

Lamin B1, an integral component of the nuclear lamina, is essential for maintaining nuclear architecture and chromatin organization. Its depletion is a recognized hallmark of cellular senescence, making it a widely used biomarker in senescence research^16^. However, quantifying perinuclear lamin B1 intensity presents significant challenges due to its specific localization at the nuclear envelope, where it resides near dense chromatin and cytoplasmic signals. SenoQuant addresses these challenges through a two-step detection approach, combining nuclear dilation and erosion to define an inner and outer boundary around the nucleus. This two-step segmentation (perinuclear rings) effectively isolates the perinuclear lamin B1 signal, reducing interference from overlapping regions while ensuring precise quantification.

To assess SenoQuant’s ability to quantify lamin B1 using its perinuclear rings feature, we analyzed lamin B1 immunofluorescence staining in sun-protected human skin from young (average age: 26.7 years) and older donors (average age: 64 years). Using the perinuclear rings feature, we quantified lamin B1 intensity in the perinuclear region and observed a decline in lamin B1 levels with age (**Figure 3a,b**).

**Figure 3.**
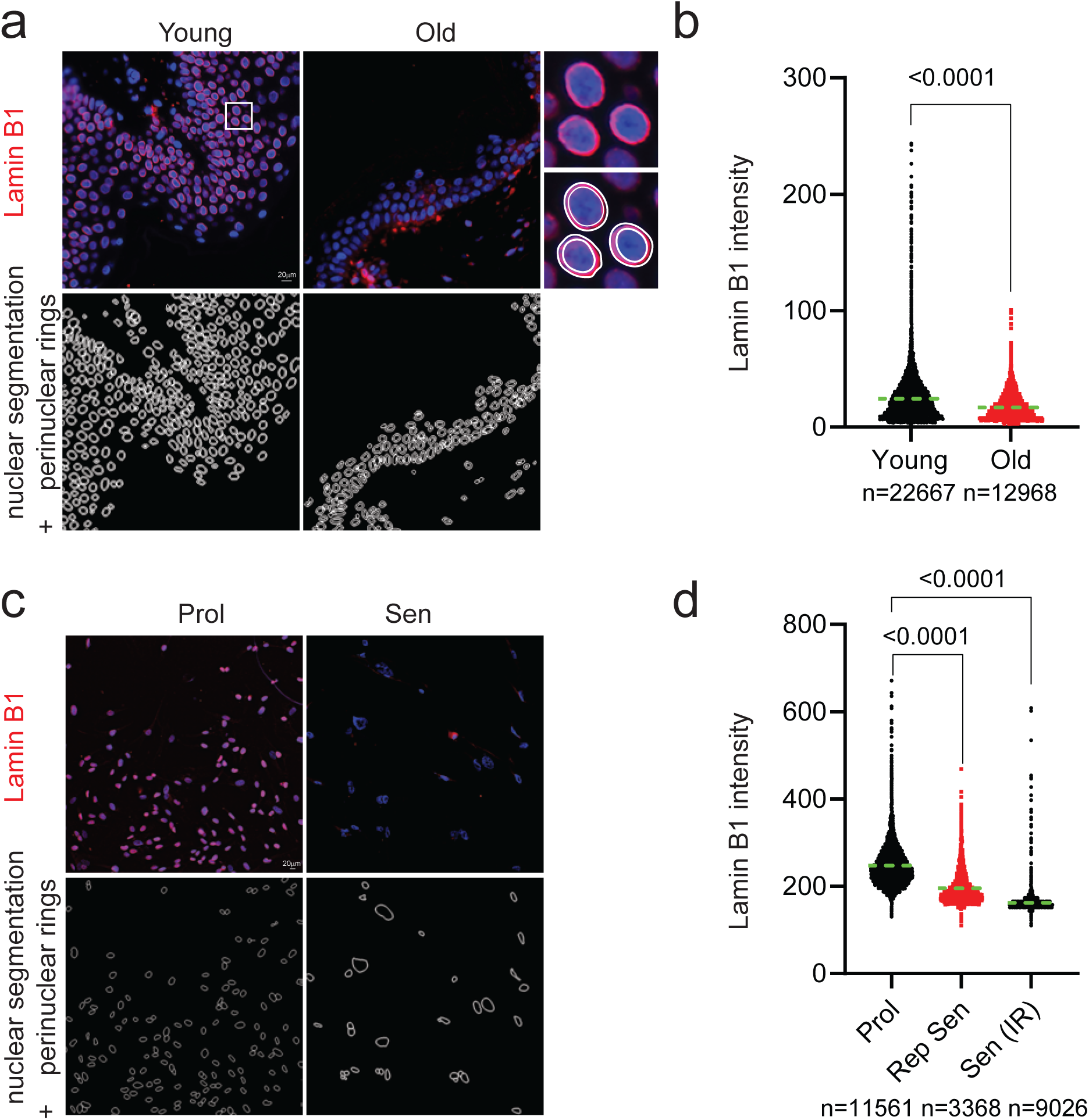
Detection of the senescence-associated marker lamin B1 using nuclear dilation. **(a)** Representative images of young and old human skin samples stained for lamin B1 (red), with nuclear membrane segmentation overlaid (double-edged, white); **(b)** Quantification of mean lamin B1 intensity in young and old human skin using SenoQuant’s nuclear segmentation and perinuclear rings features. Data are from 5 young and 5 older individuals; **(c)** Representative images of proliferative (Prol), senescent irradiated (Sen IR), and replicative senescent MRC5 fibroblasts (10 days post-irradiation) stained for lamin B1 (red), with nuclear membrane segmentation overlaid (double-edged, white); **(d)** Quantification of mean lamin B1 intensity in proliferative, replicative senescent, and irradiated senescent cells using SenoQuant’s nuclear segmentation and perinuclear rings feature. Data are shown as dot plots with the mean indicated in green. The number of cells analyzed is shown below each graph. Results are from 3–4 independent experiments. Numerical p-values are indicated in each graph (Mann-Whitney and One-Way ANOVA).

Additionally, we examined lamin B1 immunofluorescence in proliferating and senescent MRC-5 fibroblasts, where senescence was induced by either replicative exhaustion (Rep Sen) or X-ray irradiation (Sen (IR)). Our analysis revealed that lamin B1 intensity decreases in senescent fibroblasts, which is consistent with published data (**Figure 3c,d**).

### Detection of senescence-associated markers using SenoQuant’s spot detector

Telomeres, located at the ends of linear chromosomes, are protected by a group of telomere-bound proteins known as the shelterin complex^29^. This complex helps establish a looped structure, the telomere loop (T-loop), which shields chromosome ends and prevents them from being recognized as double-stranded breaks. However, once telomeres become critically short, they lose their protective cap, causing the T-loop to destabilize and exposing chromosome ends. This triggers a DNA damage response (DDR), which activates cellular senescence^30^. While telomere shortening is a key contributor to replicative senescence, recent studies show that DDR activation at telomeres can occur independently of length, both *in vitro* and in aged tissues^11,31^.

Uncapped or damaged telomeres can be identified in cells or tissues, using Immuno-FISH, which involves labeling telomeres with a Cy-3-labeled (CCCTAA) peptide nucleic acid probe and detecting DDR proteins such as γH2A.X or 53BP1. Colocalization between telomeres and DDR foci is commonly known as Telomere-associated Foci (TAF) or Telomere dysfunction induced Foci (TIF) and these are considered to be markers of cellular senescence^3^. However, manual quantification of TAF is highly time-consuming as it requires segmenting nuclei in 3D (multiple Z-stacks), detecting spot-like objects in two separate channels, and measuring individual spots and their 3D colocalization within each nucleus at high magnification (63x). The 2024 senescence detection guidelines published in Cell^3^ make it clear that TAF quantification is still technically difficult and time-consuming, and that standardized 3D colocalization is recommended. This is one of the reasons we built a dedicated spot detection module in SenoQuant, to make this analysis easier, more consistent, and less dependent on manual work.

To assess the capability of SenoQuant’s spot detector feature in quantifying TAF, we first cultured human fibroblasts until they reached replicative senescence and performed Immuno-FISH (**Figure 4a,b**). Automated quantification with SenoQuant revealed that replicative senescent cells exhibited a higher mean number of TAF compared to their proliferating counterparts. Notably, these results closely aligned with parallel manual quantification (**Figure 4c**).

**Figure 4.**
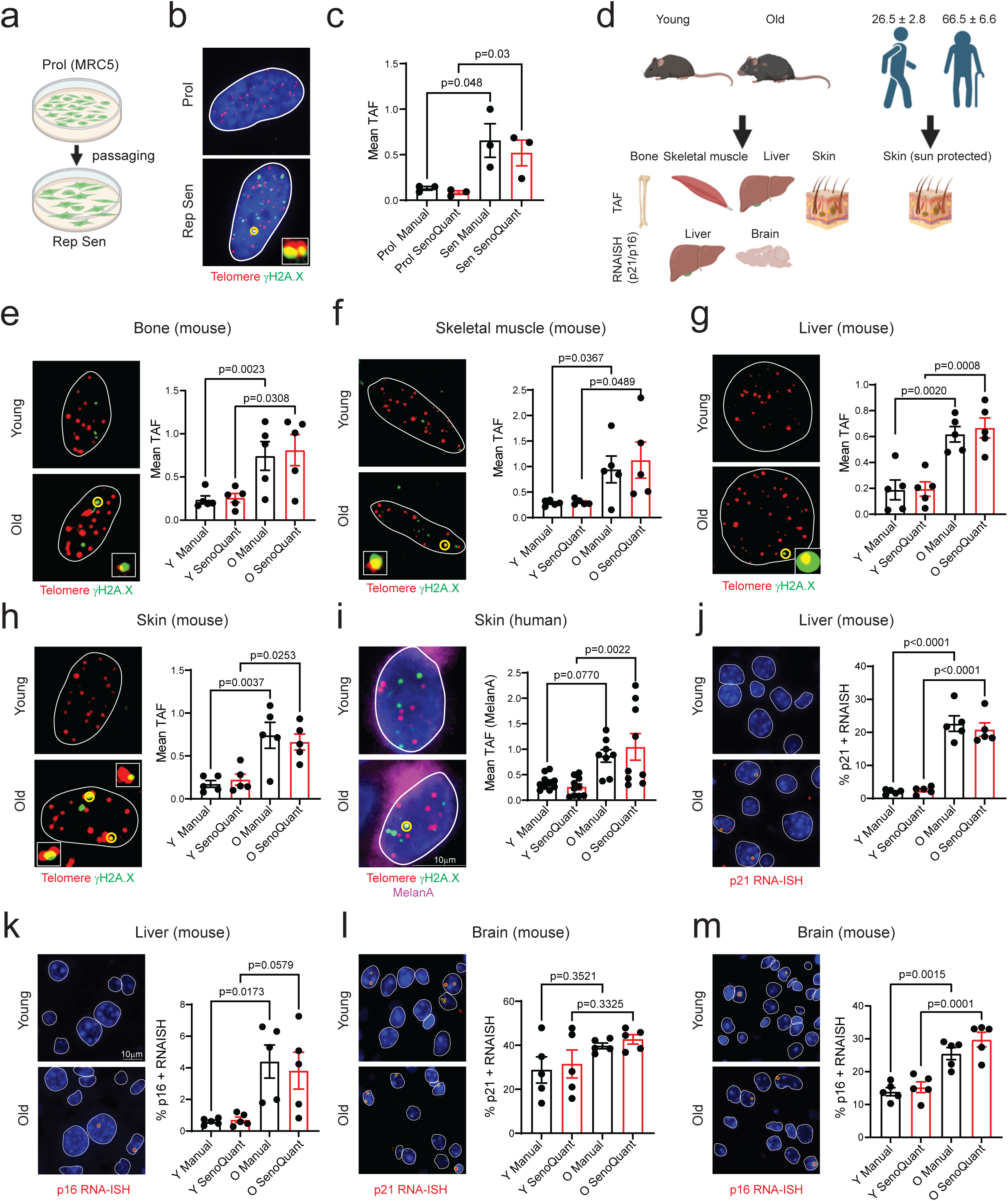
Detection of TAF and RNA-scope signals in aged tissues using SenoQuant. **(a)** Schematic representation of proliferative MRC5 fibroblasts undergoing serial passaging until reaching replicative senescence; **(b)** Representative images of TAF detection in senescent fibroblasts using fluorescence in situ hybridization (FISH) to label telomeres (red) and γH2A.X (green). Regions of 3D colocalization are circled in yellow, with nuclear segmentation outlined in white; **(c)** Quantification of mean TAF per nucleus, comparing manual annotation with SenoQuant-based automated analysis; **(d)** Experimental overview illustrating mouse and human tissue analyses across various organs using spot detection for TAF and RNA-ISH *in vivo*. Representative images of TAF detection in aged mouse tissues, including **(e)** bone (osteocytes), **(f)** skeletal muscle, **(g)** liver, and **(h)** skin, comparing young and old animals. Yellow circles indicate colocalized TAF signals. Corresponding quantification of mean TAF per nucleus is shown, comparing manual and SenoQuant-based analyses; **(i)** Representative images of TAF detection in aged human skin, with colocalization events marked by yellow circles. Melanocytes are identified using MelanA staining (purple). Quantification of mean TAF per nucleus is provided, comparing manual annotation and SenoQuant; **(j-m)** Representative images of RNA in situ hybridization (RNA-ISH) for p21 and p16 mRNA in aged mouse tissues, including liver **(j, k)** and brain **(l, m)**, comparing young and old animals. p21 RNA-ISH is shown in **(j)** and **(l)**, while p16 RNA-ISH is displayed in **(k)** and **(m)**. Yellow circles indicate detected RNA-ISH signals (red), with nuclear segmentation (white) overlaid on DAPI staining. Corresponding graphs depict the percentage of positive cells, quantified manually and using SenoQuant. Quantitative data are presented as mean ± standard error. Statistical significance is indicated numerically for each graph, with unpaired t-tests used for *in vitro* comparisons and one-way ANOVA for *in vivo* analyses. Black bars represent manual quantifications, while red bars indicate SenoQuant-based measurements. Scale bars represent 10 µm (20× and 63x objectives).

To evaluate the ability of SenoQuant’s spot detector feature to analyze TAF in tissue sections during aging, we collected liver, skin, bone, and skeletal muscle samples from young (4-6 months) and old (20-24 months) mice and performed Immuno-FISH. We then quantified TAF both manually, in a blinded manner, and using SenoQuant. Our results show that the mean number of TAF increased with age in all tissues analyzed. Importantly, the results obtained using SenoQuant closely matched those obtained through manual assessment (**Figure 4d-h**). We further tested SenoQuant’s ability to measure TAF in human tissues with age. Given that human telomeres are significantly shorter than those in mice, detecting and quantifying them using FISH-based methods can be more challenging. Previous work from our laboratory demonstrated an age-related increase in TAF in sun-protected human skin, specifically in melanocytes^32^. To build on these findings, we used SenoQuant to specifically quantify TAF in cells marked with the melanocyte marker MelanA. Our analysis revealed an age-dependent increase in the mean number of TAF in MelanA positive cells, consistent with results obtained through manual quantification (**Figure 4i**).

One of the main challenges in detecting senescence-associated markers has been the lack of specific antibodies, especially for p16 detection in murine tissues. Although recent studies have successfully validated the specificity of certain antibodies, researchers often rely on RNA *in situ* hybridization (RNA-ISH) to detect markers like p21, p16, or SASP factors at the mRNA level^28,33,34^. In this study, we collected liver and brain samples from young and old mice and performed p21 and p16 RNA-ISH^15,33,34^. Our results showed an age-related increase in both p21 and p16 expression in the liver, while in the cortex, only p16 levels significantly increased with age (**Figure 4j-m**). Notably, the results from manual quantification of RNA-ISH signals closely matched those obtained using SenoQuant, demonstrating its reliability for quantifying this data modality.

### Application of SenoQuant to spatial proteomics in aged tissues

As previously mentioned, due to the lack of specific markers, characterizing senescence benefits from omics approaches that can simultaneously analyze multiple markers. Spatial proteomics methods enable comprehensive profiling of senescence and cell-type-specific markers, as well as their spatial distribution. However, these methods require sophisticated analytical tools to manage the complexity of the data^9^.

To evaluate SenoQuant’s capability, we analyzed spatial proteomics datasets generated using either the Akoya PhenoCycler or 4i (Iterative Indirect Immunofluorescence Imaging) high-dimensional imaging platforms that use cyclic immunofluorescence. We examined four human tissues: sun-protected skin, colon and lung using different antibody panels designed to detect specific cell types and established senescence markers, including p21, p16, lamin B1, HMGB1, and SASP factors.

First, we used SenoQuant to segment all nuclei stained with DAPI. We then applied nuclear dilation and cytoplasmic segmentation to capture perinuclear or cytoplasmic markers, such as lamin B1 and secreted cytokines/chemokines. Next, we utilized the “Quantification” tab to configure a Markers feature by adding all segmentations and channels with corresponding labels. The resulting data were exported as an Excel file containing the spatial coordinates of each cell and markers per label, along with morphological parameters and fluorescence intensity measurements. These data were subsequently analyzed using R and Python (**Figure 5a**).

**Figure 5.**
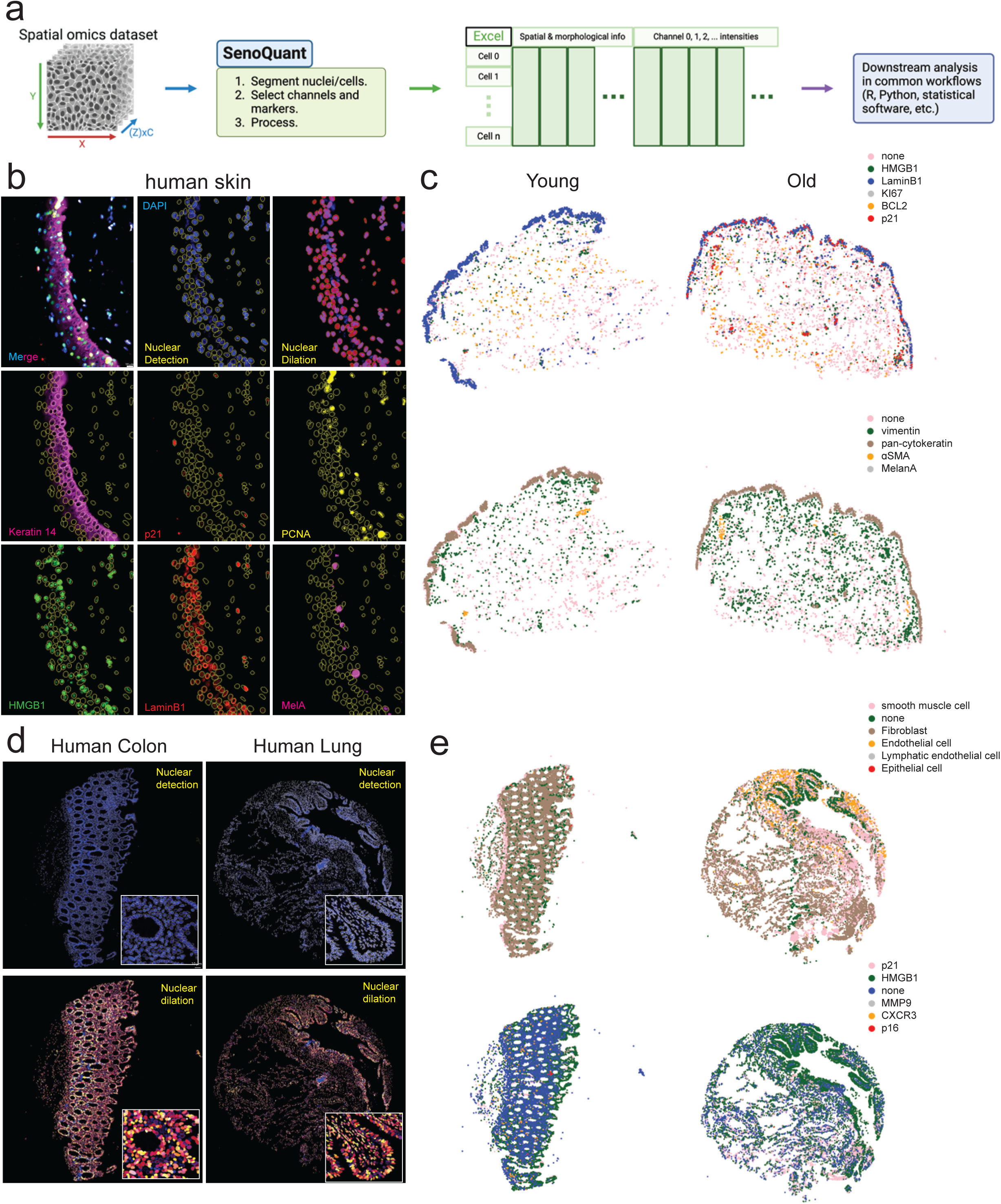
Analysis of spatial proteomics data using SenoQuant. (a) Schematic representation of the spatial proteomics analysis pipeline using SenoQuant. The workflow begins with nuclear segmentation, followed by marker selection and channel assignment based on user-defined labels. The processed data are exported as an Excel file containing all extracted measurements. Downstream analyses, including thresholding, quality control, filtering, and spatial distribution assessments, can be performed using R or Python; (b) Representative PhenoCycler images of human skin, visualizing key senescence-associated markers. The panel includes a merged image alongside individual channels: nuclear detection (yellow), nuclear dilation (red), Keratin 14 (pink), p21 (red), PCNA (yellow), HMGB1 (green), lamin B1 (red), MelanA (pink), and DAPI (blue); (c) Spatial distribution analysis of key senescent markers and cell types in young and aged human skin, highlighting differences in marker expression patterns across aging tissues; (d) Representative 4i images from human colon and lung tissue microarrays (TMA), showing nuclear detection (yellow), DAPI (blue), and nuclear dilation (multiple colors) to delineate different cell compartments; (e) Spatial distribution analysis of major cell types and senescence markers in human colon and lung tissues, demonstrating variations between different tissue types. Scale bar represents 10 µm (20× objective).

Figure 5b-e highlight PhenoCycler and 4i images illustrating nuclear detection of DAPI, nuclear dilation, and several senescence-associated markers in aged skin, colon, and lung. Following data acquisition, downstream analyses were performed to compare young and aged tissues. The spatial distribution of various cell-type and senescence-associated markers was visualized using R, with distinct colors representing different markers, providing insights into their localization and abundance (Figure 5c,e).

These results demonstrate that SenoQuant can accurately quantify the intensities for each marker based on its localization and allow us to generate maps for each tissue indicating the spatial location of each cell and marker. Importantly, it allowed us to observe the presence and spatial location of established senescence-associated markers in human tissues but also cell type specific markers (Figure 5).

### Incorporating AI-based senescence prediction into SenoQuant

A key feature of the SenoQuant platform is its ability to integrate custom AI models for senescence analysis. One emerging area in the field is the use of AI to identify senescent cells based on morphology alone, without relying on traditional biomarkers like p16 or p21. Although these morphology-based models have shown promise *in vitro*^35-38^, they often fail to achieve accurate predictions at the single-cell level in complex tissue environments^35,36^.

To address this limitation, we developed SenCeption, a proof-of-concept AI model within SenoQuant that predicts whether individual cells are p21-positive based solely on DAPI-stained nuclei in human skin tissue. For training, we used multiplexed images of DAPI and p21 obtained with the Akoya Phenocycler (formerly CODEX) from both young and old sun-protected human skin samples. The model uses convolutional neural networks (CNNs) to automatically extract morphological features from DAPI images and segmentation masks. It takes as input a DAPI-stained image of a single nucleus and its corresponding binary segmentation mask, and outputs a prediction of whether that cell is p21-positive or negative (Figure 6a).

**Figure 6.**
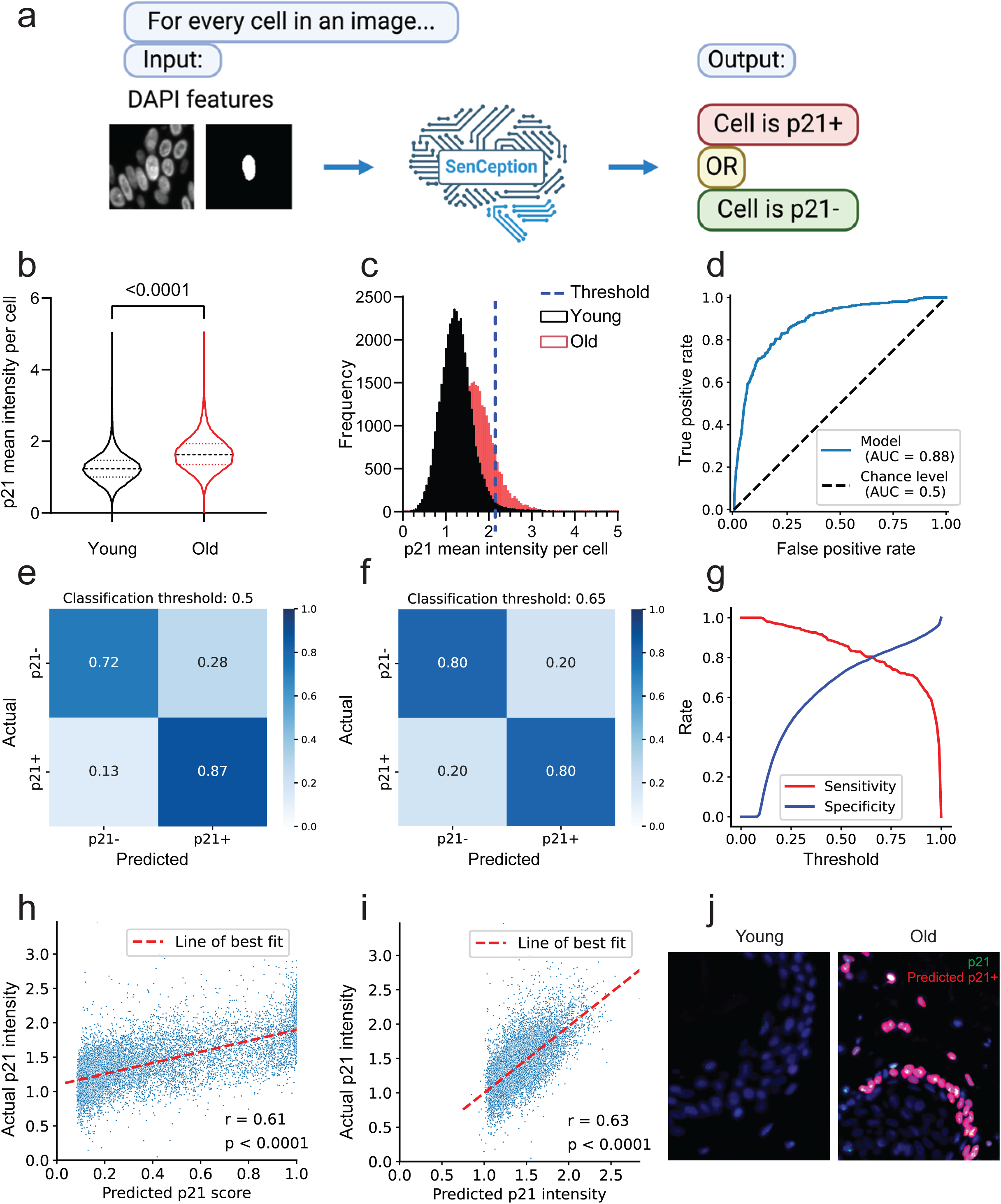
SenCeption design and performance. (a) Overview of the SenCeption model. It takes a cell’s DAPI features and predicts if the cell is p21+ or p21-; (b) Quantification of the dataset used in the development of SenCeption. For each cell, its mean intensity in the p21 channel was calculated. Then, the per-cell mean intensities were grouped by the donor’s age and compared; (c) Histogram showing the per-cell mean intensities of young and old samples and the ground truth threshold; (d) Receiver operating characteristic (ROC) curve of the classification model; (e) Confusion matrix of the classification model at classification threshold = 0.5; (f) Confusion matrix of the classification model at classification threshold = 0.65; (g) Sensitivity and specificity of the classification model at different classification thresholds; (h) Correlation between the classification model’s predicted p21 score and the actual normalized p21 intensity. Each dot is a cell in the testing set; (i) Correlation between the regression model’s predicted p21 intensity and the actual normalized p21 intensity. Each dot is a cell in the testing set; (j) Representative images of the outputs of SenCeption when integrated into SenoQuant’s AI prediction interface.

We first confirmed that p21 expression increases with age in our dataset (Figure 6b). To define ground truth for training, we set a threshold of 1.5 standard deviations above the mean of the normalized p21 intensity values, consistent with visual inspection. Using this threshold, 12.14% of cells in old samples and 2.55% in young samples were labeled as p21-positive, which aligns with previously published findings (Figure 6c)^39,40^.

We then evaluated the performance of SenCeption on an independent test set. The model achieved approximately 80% multiclass accuracy and an area under the receiver operating characteristic curve (AUROC) of 0.88 in predicting p21 status at the single-cell level (Figure 6d**,e**). A sensitivity–specificity balance of 80% can be obtained by adjusting the classification threshold from the default 0.5 to 0.65 (Figure 6f**,g**). This threshold can be fine-tuned by users when applying the model to datasets with different staining intensities or imaging conditions.

Importantly, the output scores from SenCeption are biologically meaningful. Although trained as a binary classifier (p21+ vs. p21–), the prediction scores strongly correlate with measured p21 intensity values (r = 0.61, p < 0.0001), indicating that the model captures p21 expression as a continuous variable (Figure 6h). We also trained a separate regression model to directly predict p21 intensity from DAPI features, which showed a slightly higher correlation (r = 0.63, p < 0.0001) (Figure 6i).

SenCeption predictions can be easily visualized using SenoQuant’s built-in AI prediction interface. Predicted single-cell p21 scores are overlaid on segmentation masks and displayed as a new image layer in napari, allowing users to visually assess model performance and export the data for downstream analysis (Figure 6j).

## Discussion

As members of the SenNet consortium^19^, we developed SenoQuant as a comprehensive and unbiased platform for senescence marker analysis, with a user-centric approach targeting the imaging techniques and markers frequently employed in senescence research^4,9^. The platform was intentionally designed to be accessible to investigators without programming expertise, while offering both flexibility and analytical depth for advanced research. Unlike traditional tools such as ImageJ^41^ and CellProfiler^42^, which require extensive manual adjustments and customization, SenoQuant provides a unified workflow, reducing bias, and improving reproducibility. It offers features for detecting markers at different cellular locations (nuclear/cytoplasmic), accommodating nuclei of diverse shapes, identifying spot-like objects, and integrating emerging technologies like multiplex imaging and biomarker prediction. Pre-trained AI models enable precise nuclear and cytoplasmic segmentation in human and murine tissues, as well as *in vitro* fibroblast cultures. Benchmarking comparisons confirm that SenoQuant’s nuclear segmentation models performs among the top models, with the custom default_2d being comparable to the best publicly available Cellpose variants (**Extended Data Figure 3**). Integrated spot detection and 3D colocalization allow reliable identification of Telomere Associated Foci (TAF)^11^ and RNA ISH signals, while cytoplasmic algorithms detect extraneous signals like cytokines and membrane proteins. The standardized AI prediction interface allows users to develop and test their custom models with ease. Data output is saved in CSV or XLSX format for downstream single-cell analysis, enabling studies of protein expression, spatial distribution, and cell-to-cell interactions. All settings can be saved for reproducibility and batch processing. SenoQuant is freely available as a napari plugin, supported by tutorial videos and comprehensive documentation (https://haamsree.github.io/senoquant/).

Enabled by SenoQuant’s AI prediction interface, we developed SenCeption as a proof of concept to predict the senescence-associated biomarker p21 at single-cell resolution in human skin from DAPI features alone. It represents a significant advancement in senescence-associated marker prediction in tissues compared to existing morphology-based approaches that rely on *in vitro* training data^35-38^. While previous methods have largely focused on bulk trends and struggled with single-cell accuracy *in vivo*, SenCeption achieves 80% accuracy with 0.88 AUROC at single-cell level directly in human tissue. This performance demonstrates that vision AI models trained with high-quality *in vivo* data can successfully extract meaningful senescence-associated features from minimal input information like DAPI features, dramatically reducing experimental complexity compared to traditional immunostaining methods.

While SenoQuant provides significant advantages over existing tools, several limitations remain. SenoQuant currently supports spatial proteomics technologies such as 4i and PhenoCycler but is not yet compatible with spatial transcriptomics. The alignment of imaging channels requires preprocessing using external computational pipelines before integration, and the platform’s versatility means that users are often required to adjust analysis parameters for specific experimental conditions. SenCeption, which currently focuses on p21 prediction as a proof-of-concept due to its abundant expression in skin and relevance to senescence and physiology^39,43^, is limited by its exclusive training on skin tissue and by the transient nature of p21 expression^4^, which is not sufficient for the definitive identification of senescent cells. As a result, we do not expect the model to generalize to other tissues or physiological states, and its current scope is restricted to demonstrating feasibility of tissue-based morphology prediction.

Looking forward, expanding SenoQuant’s compatibility to spatial transcriptomics and refining integration workflows for multi-channel datasets will further advance its utility. For SenCeption, future directions include incorporating additional tissues, a broader range of senescence-associated biomarkers (such as p16, lamin B1, HMGB1, and SASP factors), and input types such as H&E, with the goal of enabling stain-free senescence detection at single-cell resolution. These enhancements would increase throughput, reduce costs, and accelerate research and translational applications in aging and age-related diseases.

## Materials and Methods

### Development of the segmentation models Datasets

#### Datasets used to train the default_2d model

1. Manually annotated images generated from our laboratory.
2. A subset of the Data Science Bowl 2018 (https://www.kaggle.com/c/data-science-bowl-2018) dataset.
3. A subset of S-BSST265 (https://www.ebi.ac.uk/biostudies/bioimages/studies/S-BSST265).
4. TissueNet v1.1 (https://datasets.deepcell.org/data) training and validation subsets.

The combined dataset contains 5837 annotated images. 2691 and 3146 images were assigned to training and validation, respectively.

#### Datasets used to train the default_3d model

1. Manually annotated images generated from our laboratory.
2. Synthetic images generated with various perturbations (sparse, dense, shapes, sizes, etc.).

The combined dataset contains 98 images (53 annotated, 45 synthetic). 83 and 15 images were assigned to training and validation, respectively.

The images mentioned in the datasets above include *in vitro* and *in vivo* samples from various human and murine tissues (skin, bone, liver, brain, etc.). Manual annotations were performed using Labkit in Fiji in 2D and 3D following the instructions from https://github.com/stardist/stardist.

### Benchmarking

The model benchmarking in **Extended Data Figure 3** was performed using the TissueNet v1.1 testing set. default_2d and cpsam are included in the current release of SenoQuant. cp_cyto3 and cp_nuclei are Cellpose 3 default models. The object diameter parameter of each model was set to 16.18 pixels, which is the mean object diameter of the testing set. Details regarding the benchmarking can be found in the model_benchmarking folder of the project (https://github.com/HaamsRee/senoquant/tree/main/model_benchmarking).

### Model details

Details about the implemented models can be found at https://github.com/HaamsRee/senoquant and https://haamsree.github.io/senoquant.

### Computer setup and specifications

Training was performed using a server with two Intel Xeon E5-2620 v4 CPUs and 256GB RAM with two NVIDIA GV100 [TITAN V] GPUs.

### Development of SenCeption

#### Dataset preparation and quantification

p21 and DAPI channels of the ten PhenoCycler skin images from ten different young and old donors were used to create the dataset. Each image was first processed to have the same float32 number format and 0.25 μm physical pixel dimensions. Then, the white tophat algorithm with a disk kernel with a 30-pixel radius was used to subtract the background. The background-subtracted DAPI images were segmented with a fine-tuned Cellpose3 model to obtain the nuclear masks^44^. The masks were then manually checked for quality and any that overlaps with autofluorescence were discarded, which left us with ∼74,000 cells.

The images were obtained on three separate days, with eight samples imaged on the first two and two samples (one young 24 y/o female and one old 85 y/o male) imaged on the last. To normalize the images, we grouped them into three batches accordingly and calculated batch intensity medians of the nuclei identified by segmentation for every batch and every channel. Images in the same batch were then divided by the median intensities of the batch. The median-divided images were further transformed by the inverse hyperbolic sine transform with a cofactor of 0.5 to reduce the effects of outliers.

For every nucleus identified, we calculated the mean intensity of its p21 channel as its p21 expression. Based on visual assessment and existing literature, we set the labeling threshold at 1.5 standard deviations from the dataset mean, which resulted in 12.14% of the cells in the old samples and 2.55% of the cells in the young samples labeled as p21+. Using segmentation mask centroids, for each cell, we prepared two 256x256 layers as model inputs: The cell’s DAPI image and binary nuclear segmentation mask, both with the cell of interest centered. We then split the dataset into training, validation, and testing sets. We isolated cells from the two samples imaged on the last day as the independent testing set and grouped the rest into training (80%) and validation (20%) sets.

#### Model architecture

We tested a variety of CNN backbones and used a modified ResNet-50-CBAM^45^ that performed the best. SenCeption accepts 2x256x256 inputs and outputs a two-dimensional logit vector. After Softmax activation, the values of the first and second elements add up to one can be interpreted as p21- and p21+ scores, respectively. The higher the p21+ score (and the lower the p21- score), the more the model predicts the cell at its input is p21+. The default binary classification threshold was set at 0.5. The regression model takes the same 2x256x256 inputs and outputs a single value that corresponds to the predicted p21 intensity of the cell at its input. PyTorch and PyTorch Lightning were used to build the model^46,47^.

#### Model training and testing

Ray Tune was used to identify the hyperparameters, including input rescaling, sample weights, learning rate, weight decay, gradient accumulation steps, and data augmentation, that resulted in the best performing models^48^. Each experiment had a time budget of two days and the goal of maximizing validation multi-class accuracy (mean of accuracies of each class). For the regression model, the goal was to minimize validation mean squared error (MSE). We used the Async Successive Halving scheduler and the Optuna (TPESampler) search algorithm. AdamW was used as the optimizer. The best models were tested with the independent testing set and metrics including multi-class accuracy, AUROC, and Pearson correlation coefficient were calculated.

SenCeption experiments were performed on a workstation with an Intel Xeon W7-3565X CPU, 256GB RAM, and four NVIDIA RTX 6000 Ada 48GB GPUs.

##### Experimental procedures

#### Tissue culture

Human embryonic lung MRC5 fibroblasts (ATCC) were grown in Dulbecco’s modified Eagle’s medium (Sigma-Aldrich, D5796) supplemented with 10% heat-inactivated fetal bovine serum (FBS), 100LULml−1 penicillin, 100LμgLml−1 streptomycin and 2 mM L-glutamine and maintained at 37 °C under 5% CO_2_. MRC5 fibroblasts were cultured in atmospheric oxygen conditions.

#### Stress-induced senescence

Cells were exposed to 20Gy X-ray irradiation. Cells were cultured for another 10 days after irradiation, then coverslips were fixed using 2% PFA for 5 min, then stored either in the fridge at 4 degrees or at -80 degrees for long term storage. Senescence was confirmed by the presence of multiple established markers such as p16, p21, γH2A.X and loss of lamin B1 expression by immunostaining.

#### Animal experiments

All animal experiments adhered to protocols approved by the Mayo Clinic Institutional Animal Care and Use Committee (IACUC), unless otherwise stated. The study employed two distinct age groups of C57BL/6 mice: young mice (4-6 months) and aged mice (20–22 months): These mice were maintained in a pathogen-free facility. They were housed in same-sex cages (3–5 mice per cage) on a 12-hour light/dark cycle at 23–24°C with ad libitum access to regular chow and water.

Euthanasia was performed for mice meeting pre-defined humane endpoints. Investigators remained blinded to group allocation throughout the experiments and outcome assessments. Data collection and analysis were also conducted in a blinded manner to minimize potential bias.

### Analysis of telomere-associated foci and lamin B1 in human sun-protected skin

Archived micrographs of immunofluorescence or Immuno-FISH in human sun protected skin from ^32^ were either reanalyzed manually or using SenoQuant.

### Immunocytochemistry

Fibroblasts grown on coverslips were fixed using 2% paraformaldehyde in PBS for 10Lmin. Cells were washed in PBS (x2; 5min each) and then permeabilized in PBG-Triton (PBS, 0.4% fish skin gelatin, 0.5% BSA, 0.5% Triton X-100) for 45 min. Subsequently, cells were incubated with primary antibody overnight at 4 °C. After PBS washes, secondary antibodies were applied and incubated for 60 min at room temperature. Coverslips were mounted onto superfrost glass microscope slides with ProLong Gold Antifade Mountant with DAPI (Invitrogen). A list of the antibodies used is provided in **Table 2**.

### Formalin-fixed paraffin-embedded Immuno-FISH

Tissue sections were deparaffinized in 100% Histoclear, then hydrated using a graded ethanol series of 100, 90 and 70% ethanol (twice for 5 min each) and washed twice for 5Lmin in distilled water. Antigen retrieval was performed in 0.01 M citrate buffer (pHL6.0) and heated in a microwave until boiling for 10 min. The sections were allowed to cool to room temperature and then washed in distilled water for 5 min followed by 5 min in PBS. Blocking was then performed using normal goat serum (1:60) in BSA/PBS for 30 min followed by overnight incubation with rabbit monoclonal anti-γH2AX antibody (1:400, 9718; Cell Signaling) at 4 °C. After three PBS washes, tissues were incubated with a goat anti-rabbit biotinylated secondary antibody (1:200, PK-6101; Vector Labs) for 30 min at room temperature followed by three washes in PBS and incubated with Streptavidin cy5 (1:500, (SA-1500-1); Vector Labs) for 30Lmin at room temperature. The tissues were then washed three times in PBS and incubated in 4% paraformaldehyde in PBS for 20 min for cross-linking. Then, after three PBS washes, sections were dehydrated in graded cold ethanol solutions (70, 90, 100%) for 3 min each and were then allowed to air dry. Next, 10 μl of PNA hybridization mix (70% deionized formamide (Sigma-Aldrich), 20LmM MgCl2, 1LM Tris pHL7.2, 5% blocking reagent (Roche) containing 2.5Lμg ml–1 Cy-3-labelled telomere-specific (CCCTAA) peptide nucleic acid probe (PANAGENE) was added to sections and denaturation was done for 10 min at 80 °C. Sections were then incubated in PNA hybridization mix for 2 h at room temperature in the dark to allow hybridization. Tissues were then washed in 70% formamide in 2× SSC for 10Lmin, followed by one wash in 2× SSC for 10 minutes and a PBS wash for 10 min. Tissues were mounted using ProLong Gold Antifade Mountant with DAPI (Invitrogen) and imaged using in-depth z stacking (a minimum of 40 optical slices with a 63x objective).

### RNA in situ hybridization (RNA-ISH)

RNA-ISH was performed per the RNAScope protocol from Advanced Cell Diagnostics Inc. (ACD): RNAScope Multiplex Fluorescent Assay v2. Briefly, the assay allows simultaneous visualization of up to 4 RNA targets, with each probe assigned to a different channel (C1, C2, or C3 or C4). p21 (CDKN1A) signal amplified using the HRP-C1 linked with a secondary fluorophore, Opal 570 (detected in the Cy3 range), followed by a subsequent detection of p16/p19 (CDKN2A) –c3, the signal is amplified by the HRP-C3 linked with a secondary fluorophore, Opal 650 (detected in the Cy5 range). Tissue sections were then mounted using ProLong Gold Antifade Mountant with DAPI (Invitrogen). Sections were imaged using single Z plane. RNA-ISH-positive cells on tissue were quantified normalized against total cells on the tissue surface.

### Microscopy

Imaging was performed using a Leica SP8 widefield microscope using various objectives (x20, 40x, 63x) depending on experiment requirements.

Visualization and manual analysis were performed using FIJI or napari viewer with SenoQuant plugin.

### Statistical analysis

GraphPad Prism v.10.02 was used for statistical analysis; the results were statistically significant when PL≤L0.05. For normally distributed data, the differences between two groups were tested for statistical significance using an independent-sample two-tailed t-test. For data that were normally distributed and when there was more than one group, one-way ANOVA was used, with Tukey’s comparison post hoc test.

### Ethics statement

All animal experiments were performed according to protocols approved by the Institutional Animal Care and Use Committee (IACUC) at Mayo Clinic.

The Tissue Macro Array (TMA) was constructed in the Pathology Research Core of Mayo Clinic under the IRB - 14-006858 (Control and Assay Development Tissues for use in the Pathology Research Core-II).

**Table 1:**
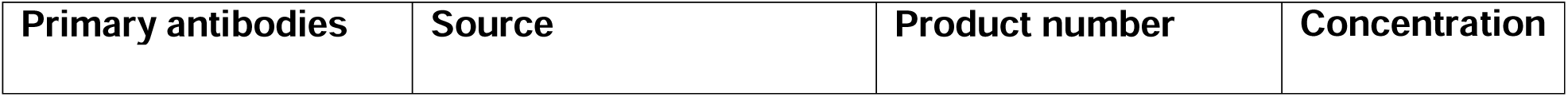
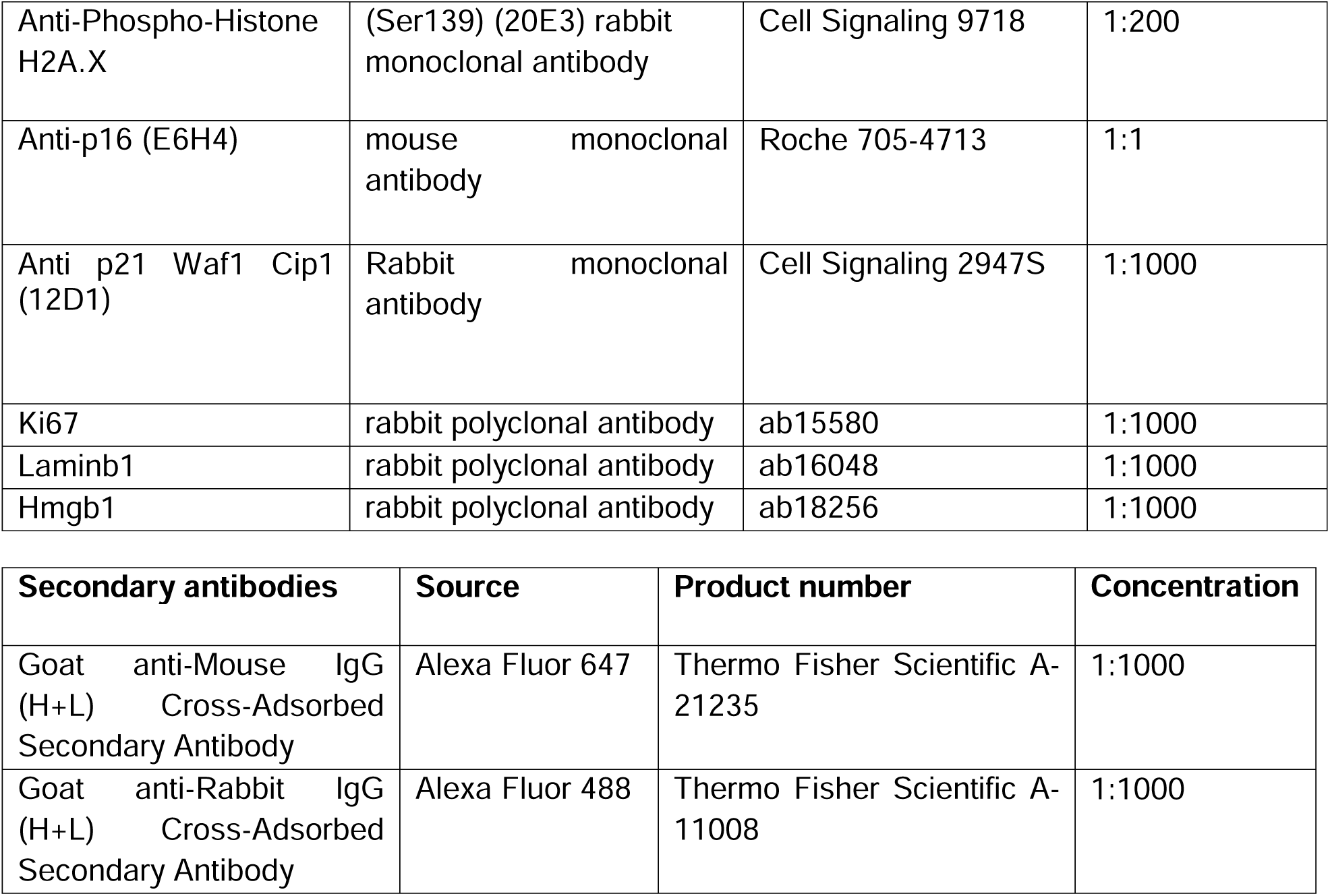
Antibodies list used for conventional immunofluorescence.

| Primary antibodies | Source | Product number | Concentration |
| --- | --- | --- | --- |
| Anti-Phospho-Histone H2A.X | (Ser139) (20E3) rabbit monoclonal antibody | Cell Signaling 9718 | 1:200 |
| Anti-p16 (E6H4) | mouse monoclonal antibody | Roche 705-4713 | 1:1 |
| Anti p21 Waf1 Cip1 (12D1) | Rabbit monoclonal antibody | Cell Signaling 2947S | 1:1000 |
| Ki67 | rabbit polyclonal antibody | ab15580 | 1:1000 |
| Laminb1 | rabbit polyclonal antibody | ab16048 | 1:1000 |
| Hmgb1 | rabbit polyclonal antibody | ab18256 | 1:1000 |

| Secondary antibodies | Source | Product number | Concentration |
| --- | --- | --- | --- |
| Goat anti-Mouse IgG (H+L) Cross-Adsorbed Secondary Antibody | Alexa Fluor 647 | Thermo Fisher Scientific A-21235 | 1:1000 |
| Goat anti-Rabbit IgG (H+L) Cross-Adsorbed Secondary Antibody | Alexa Fluor 488 | Thermo Fisher Scientific A-11008 | 1:1000 |

### PhenoCycler-Fusion imaging of skin sections

**Table 2:** PhenoCycler sample information.

| ID | Group | Age | Sex | SenCeption train/test |
| --- | --- | --- | --- | --- |
| 0 | Old | 65 | Male | Train |
| 1 | Old | 67 | Female | Train |
| 2 | Old | 82 | Female | Train |
| 3 | Old | 83 | Male | Train |
| 4 | Old | 85 | Male | Test |
| 5 | Young | 28 | Female | Train |
| 6 | Young | 23 | Female | Train |
| 7 | Young | 20 | Male | Train |
| 8 | Young | 31 | Male | Train |
| 9 | Young | 24 | Female | Test |

Skin sections (**Table 2**), 5 µm thick, were cut onto standard microscope slides (Leica, cat# 3800080) and air-dried at room temperature for 48 hours before staining. Staining was performed using the PhenoCycler-Fusion (formerly CODEX) system following the vendor-provided protocol (Version 2.1.0, Akoya Biosciences) with minor modifications.

Prior to staining, tissue sections were baked at 65°C for 2 hours, deparaffinized in xylene, and sequentially rehydrated in graded ethanol solutions. Heat-induced epitope retrieval (HIER) was performed in a pressure cooker (InstantPot Duo) at high pressure for 20 minutes using Tris-EDTA pH 9 buffer (Akoya, cat# AR900250ML). After retrieval, slides were cooled at room temperature for 30 minutes, followed by a “waterfall” wash with deionized water to gently exchange the retrieval buffer and remove residual reagents. While the slides were cooling, the antibody panel was assembled by combining the staining buffer, kit-provided blockers, and 40 antibodies at their designated concentrations (**Table 3**). Tissue sections were then equilibrated in staining buffer for 30 minutes before incubation with the antibody cocktail for 3 hours at room temperature. Following incubation, the slides underwent a series of washes and three separate fixation steps to stabilize antibody binding.

To minimize background autofluorescence associated with specific tissue types and formalin-fixed paraffin-embedded (FFPE) samples, an autofluorescence quenching step was incorporated using two broad-spectrum LED light sources and a quenching solution containing 4.5% (w/v) hydrogen peroxide and 20 mM sodium hydroxide in PBS. After quenching, samples were thoroughly washed, placed in storage buffer, and stored at 4°C until imaging.

Before imaging, slides were equilibrated to room temperature, and the fluidic cell was assembled according to Akoya’s protocol. Imaging was performed using the PhenoCycler-Fusion (Version 2.1.0) at 20× magnification across 24 imaging cycles. Of these, 21 cycles contained antibody targets, one included irrelevant reporters for downstream analysis, and two were blank cycles for background subtraction and tile registration. Exposure times were set at 2 milliseconds for DAPI, 150 milliseconds for AlexaFluor™ 647, 150 milliseconds for Atto550, and 150 milliseconds for AlexaFluor™ 750.

Raw imaging data were processed using CODEX Uploader to perform stitching, drift correction, deconvolution, and cycle concatenation. The processed data were then loaded into SenoQuant for further quantification and analysis.

**Table 3:** PhenoCycler-Fusion antibodies list.

| Target | Clone | Primary Catalogue # | Primary Vendor | BX Conjugated | Channel Acquired | Custom Conjugation |
| --- | --- | --- | --- | --- | --- | --- |
| **Keratin 14** | Poly19053 | AKOYA | AKOYA | 2 | Atto550 | N |
| **p21 Waf1/Cip1** | 12D1 | 19399SF | CST | 6 | AF647 | Mayo Hybridoma core |
| **Ki67** | B56 | AKOYA | AKOYA | 47 | Atto550 | N |
| **HMGB1** | EPR3507 | AB216986 | Abcam | 135 | AF647 | Mayo Hybridoma core |
| **Lamin B1** | EPR22165-121 | AB239399 | Abcam | 28 | AF647 | Mayo Hybridoma core |
| **PCNA** | PC10 |  | Abcam | 36 | AF647 | N |
| **BCL2** | EPR17509 | AKOYA | AKOYA | 85 | Atto550 | N |
| **Vimentin** | 091D3 | AKOYA | AKOYA | 22 | AF750 | N |
| **Pan-** | AE-1/AE-3 | AKOYA | AKOYA | 19 | AF750 | N |
| **Cytokeratin** |  |  |  |  |  |  |
| **SMA** | 1A4 | AKOYA | AKOYA | 13 | AF750 | N |

### Iterative Indirect Immunofluorescence Imaging (4i)

Formalin-fixed, paraffin-embedded (FFPE) tissue microarrays (TMAs), including 5-µm-thick sections of lung and colon tissue from a 62-year-old male and an additional colon sample, were used for imaging. Sections were first deparaffinized by immersing them twice in HistoClear for 5 minutes each. This was followed by progressive rehydration through a graded ethanol series, including two washes in 100% ethanol for 5 minutes each, followed by 90% and 70% ethanol washes for 5 minutes each. After rehydration, sections were rinsed twice with distilled water for 5 minutes per wash.

Antigen retrieval was performed by immersing the sections in 0.01 M citrate buffer (pH 6.0) and heating them to boiling for 10 minutes. Following retrieval, sections were allowed to cool to room temperature and washed for 5 minutes in distilled water. To block non-specific binding, sections were incubated for 1 hour at room temperature in a blocking solution containing 1% bovine serum albumin (BSA) and 150 mM maleimide in phosphate-buffered saline (PBS). After blocking, sections were washed twice with PBS.

For antibody staining, sections were incubated overnight at 4°C with a primary antibody cocktail (see **Table 4**), diluted in blocking buffer (1% BSA in PBS). The next day, sections were washed twice in PBS before incubation with secondary antibodies for 1 hour at room temperature, followed by another two PBS washes. Nuclear staining was performed using DAPI, diluted 1:1000 in PBS, for 10 minutes at room temperature, followed by two additional PBS washes.

Mounting was performed by adding 50 µL of imaging buffer, which was prepared by dissolving 1140 mg of N-acetylcysteine in 3.5 mL of water and 6.5 mL of 1 M sodium hydroxide, with the pH adjusted to 7.5. A coverslip was carefully placed over the sections, ensuring even distribution of the mounting medium.

Imaging was conducted using an epifluorescence microscope, capturing tiled images of the entire section. After imaging, antibody elution was performed to enable subsequent rounds of staining. Sections were treated with an elution buffer containing 0.5 M glycine, 3 M urea, 3 M guanidine hydrochloride, and 70 mM tris(2-carboxyethyl) phosphine (TCEP) in water, adjusted to pH 2.5. This treatment was repeated four times, each lasting 10 minutes at room temperature with agitation. After elution, the protocol was restarted from the blocking step, with a 20-minute incubation at room temperature, to allow additional rounds of antibody staining and imaging.

**Table 4:** Antibodies used for Iterative Indirect Immunofluorescence Imaging.

| Target | Species | Brand/reference | Dilution factor |
| --- | --- | --- | --- |
| p16 | mouse | VENTANA (725-4793) | As provided by kit |
| p21 | rabbit | Cell signalling (2947S) | 100 |
| PCNA | mouse | Abcam (ab29) | 500 |
| HMGB1 | rabbit | Abcam (ab18256) | 200 |
| MMP9 | rabbit | Cell signalling (2270) | 200 |
| CXCR3 | mouse | Abcam (ab64714) | 200 |
| SMA | goat | Abcam (Ab21027) | 200 |
| IP-10 | rabbit | Abcam (ab9807) | 100 |
| e-Cadherin | <u>mouse</u> | Abcam (Ab76055) | 200 |
| Lyve1 | goat | R&D AF2089 | 200 |
| Vimentin | rabbit | Abcam (ab92547) | 100 |

### Analysis pipeline of 4i and PhenoCycler images with SenoQuant

The analysis of 4i and PhenoCycler aligned images was performed using SenoQuant, following a structured pipeline to ensure accurate quantification of cellular markers.

The process began with nuclear segmentation, where individual nuclei were identified and labeled to create a reference mask. A quality control step was then applied to each analyzed channel to exclude nuclei overlapping with regions exhibiting high autofluorescence or non-specific binding, as well as areas with folded tissues, which could introduce artifacts.

Once the nuclear labels were refined, a nuclear dilation step was applied, expanding each nucleus by 3 pixels to generate a surrounding disk that captures cytoplasmic staining. The dilation size was optimized empirically to ensure sufficient cytoplasmic coverage while minimizing overlap with neighboring cells, as this parameter varies depending on tissue and cell type. Intensity measurements were then recorded separately for both the original nuclear mask and the expanded cytoplasmic region, with user-defined settings for each channel.

Following intensity extraction, an Excel file was generated, compiling individual marker intensities and additional cellular properties for each nucleus. The next step involved defining intensity thresholds to determine marker positivity, with threshold values applied to classify each cell as positive or negative for specific markers. Positive cells were annotated within the Excel data analysis sheet for further downstream analysis.

To enhance accuracy, the dataset was manually curated and cleaned, ensuring the removal of false positives. Finally, marker expression values were transformed into binary values (0,1) to categorize cells as either positive or negative for each marker, allowing for robust classification and further quantitative assessments.

### Data analysis using R and Python

After performing image analysis on SenoQuant as well as manual quality control and thresholding, the raw counts for each marker for each cell, along with spatial distribution information based on the x- and y-values derived from SenoQuant, were read into R, where each sample was normalized and scaled in Seurat (v5.0)^49^. The samples were integrated into a single Seurat object and visualized using Seurat (v5.0) and ggplot2 (v3.5.1).

The quantifications, image processing, statistics, and visualization involved in SenCeption development were performed with Python 3 using numpy, cupy, pandas, scipy, scikit-image, cucim (https://github.com/rapidsai/cucim), bioio (https://github.com/bioio-devs/bioio), matplotlib, and seaborn^50-56^.

## Acknowledgements

This work was funded by NIH grants UH3CA268103 (JFP); R01AG068048 (JFP); R01AG82708 (JFP),); P01 AG062413 (DJ, JFP), UH3CA268202 (NN), The Glenn Foundation For Medical Research (JFP), R01 AG068182 (DJ), Hevolution/AFAR (DJ), R01 HL158532 (YSP), R01 HL171915 (YSP) NIH R03-AG082919-01 (SW), Hevolution HF-GRO-23-1199252-26, P01 AG062413 (SK), R01AG086085 (PI) (SK), R01 AG076515 (SK), U54 AG079754 (SK), Hevolution HR-GRO-23-1199144-8 (SK), The National Science Foundation Graduate Research Fellowships Program (DGE-1842487) (AJH), R01 DK128552 (JNF). While SenoQuant was funded by SenNet Common Fund award UH3CA268103, SenCeption was independently funded by Mayo Clinic.

## Author contributions

ABL and YL developed SenoQuant with contributions from JFP, JC, CN, DHIII, ACF, YH, HM, SV, MS, NN. GL. YL developed SenCeption. DGC, DS, AJH, LSG, JNF, SW and LK performed individual experiments and/or analyzed data. SW and YSP provided human tissues. SK and DJ supervised components of the study. JFP, ABL, and YL conceptualized SenoQuant and SenCeption and wrote the manuscript with contributions from all authors.

## Data availability

Data are available from the authors upon reasonable request.

## Code availability

The source code for SenoQuant is available at https://github.com/HaamsRee/senoquant. Detailed documentation and tutorials are available at https://haamsree.github.io/senoquant.

## Extended Data Figure legends

**Extended Data Figure 1.**
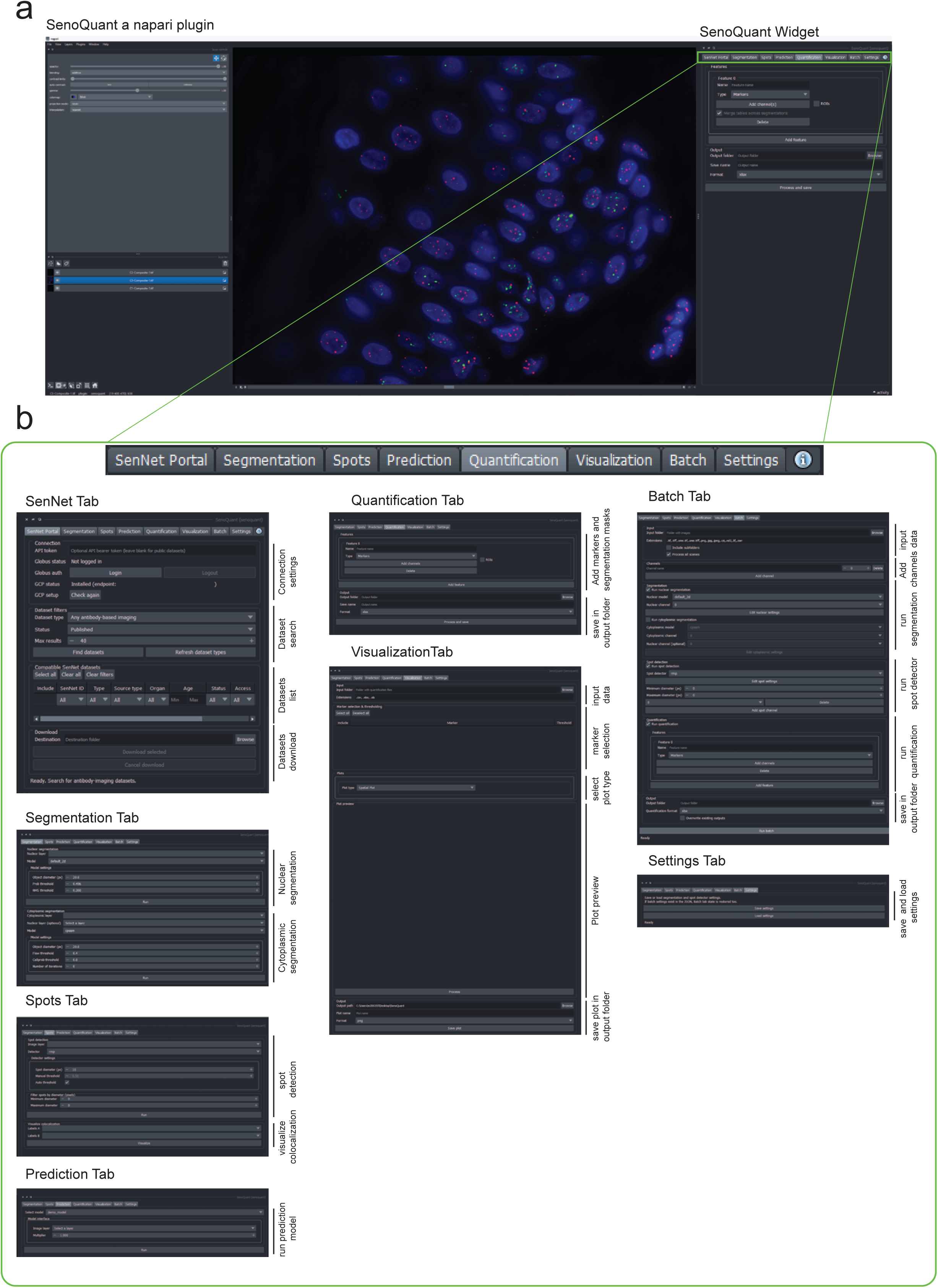
SenoQuant overview. (a) Snapshot showing the SenoQuant GUI. The green rectangle shows a zoomed version of all tabs which are displayed and annotated in detail (b).

**Extended Data Figure 2.**
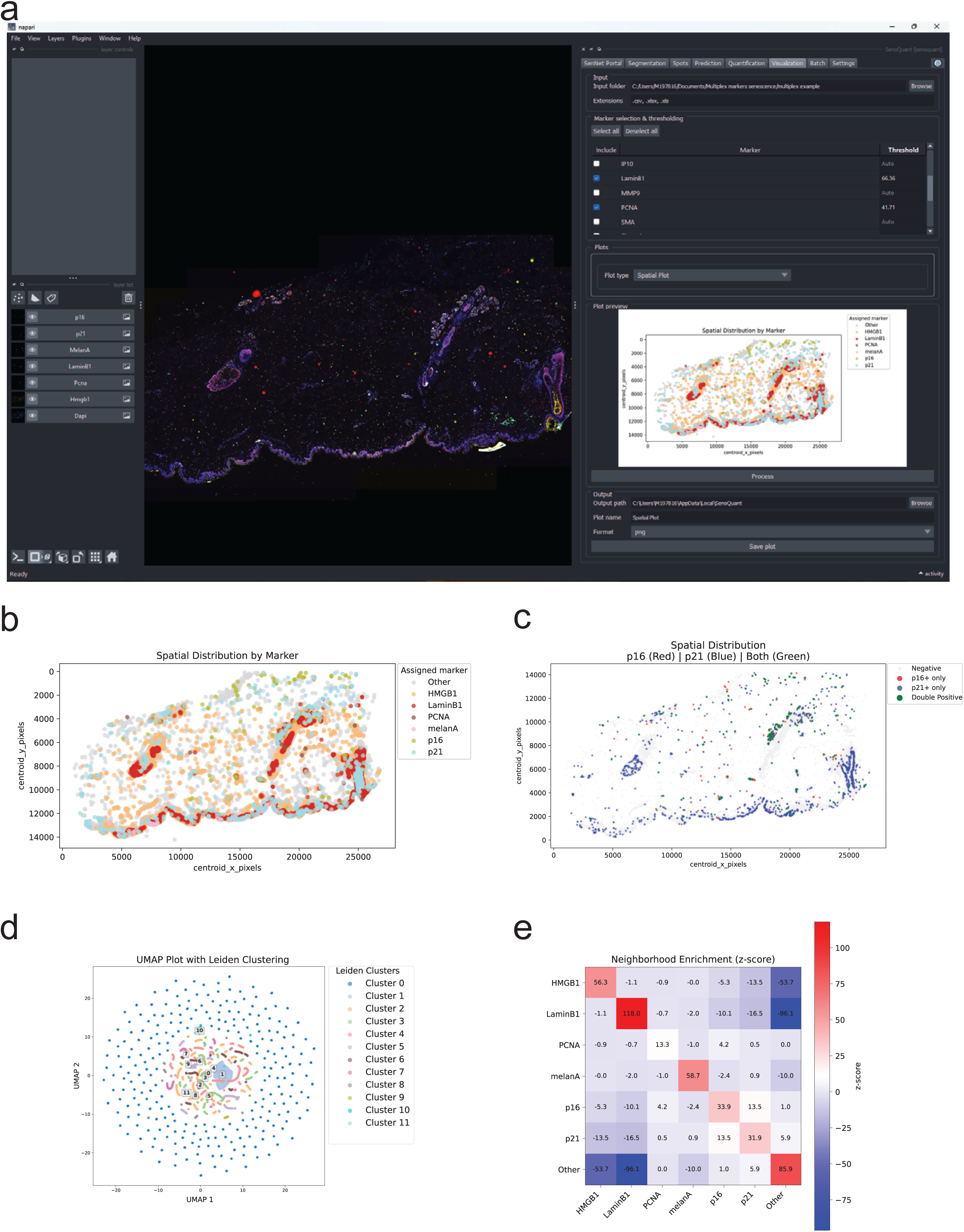
Visualization tab. (a) Snapshot showing the visualization tab using an xlsx file outputted from SenoQuant. The table in the interface has the option to include/exclude markers and indicate a positivity threshold per marker. The display below the table shows a spatial distribution plot by marker in human skin. (b-e) Examples of plots available: spatial distribution plot, spatial distribution plot using two markers counting single and double positives, UMAP plot with Leiden clustering, and neighborhood enrichment plot (z-score).

**Extended Data Figure 3.**
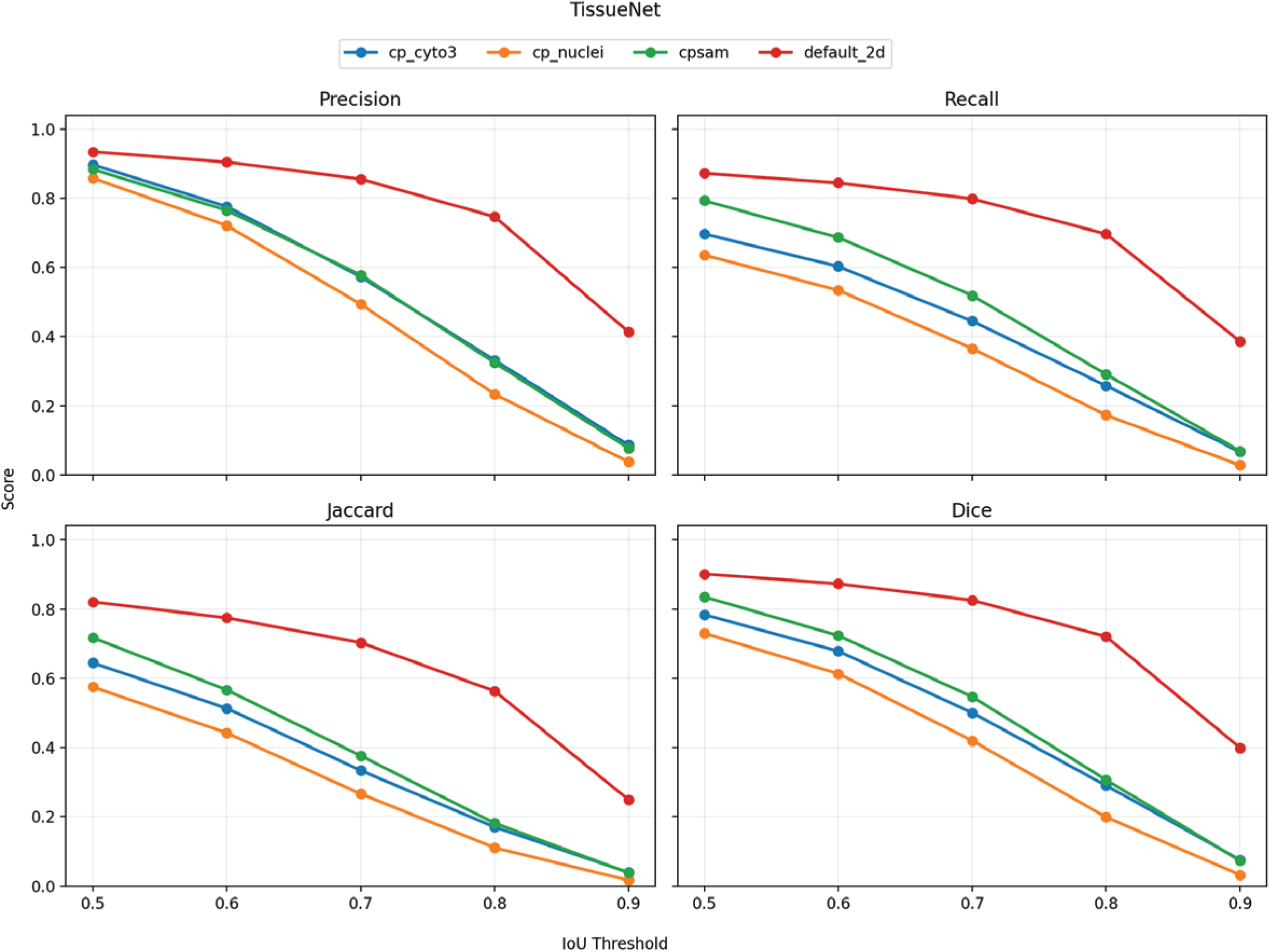
Benchmarking of nuclear segmentation models. Panels showing the precision, recall, Jaccard index, and Dice coefficient scores for a range of segmentation models evaluated at various IoUs using the TissueNet v1.1 testing set. default_2d and cpsam (Cellpose-SAM) are available in SenoQuant. cp_cyto3 and cp_nuclei are Cellpose 3 default models. The custom default_2d model in SenoQuant (red) performs among the top models across all metrics and is comparable to the best publicly available Cellpose variants.

